# Male and Female Mice Are Similarly Susceptible to Chronic Nondiscriminatory Social Defeat Stress Despite Differences in Attack Frequency from Aggressor

**DOI:** 10.1101/2025.03.25.645316

**Authors:** Allyson Bazer, Katherine Denney, Maria Chacona, Catherine Montgomery, Shriya Vinod, Urboshi Datta, Benjamin Adam Samuels

**Author notes:** **Correspondence:** Benjamin A Samuels.

## Abstract

**Rationale:** Mood disorders are often precipitated by chronic stress and can result in an inability to adapt to the environment and increased vulnerability to challenging experiences. While diagnoses of mood disorders are diagnosed twice as frequently in women than in men, most preclinical chronic social defeat stress mouse models exclude females due to decreased aggression toward female intruders.

**Objectives:** We previously reported that the chronic non-discriminatory social defeat stress (CNSDS) paradigm is effective in both sexes, allowing for comparisons between male and female mice. We aimed to improve the screening protocol to identify CD-1 aggressors for use in CNSDS and the method for determining susceptibility to CNSDS. Finally, we aimed to determine whether susceptibility to CNSDS correlated with impaired performance in a satiety-based outcome devaluation task.

**Methods:** We analyzed CNSDS screening and social defeat sessions to determine appropriate parameters for selecting CD-1 aggressors and investigated aggressions toward male and female intruder mice. We also investigated CNSDS effects on a reward valuation task.

**Results:** We observed that despite receiving fewer attacks, female mice are equally susceptible to CNSDS as males and that CNSDS abolished satiety-based outcome devaluation in susceptible male and female mice, but not in resilient male and female mice.

**Conclusions:** These data suggest that CNSDS-defined susceptible and resilient phenotypes extend to reward behaviors.

## INTRODUCTION

Millions of people suffer from major depressive disorder (MDD), a mood disorder that the World Health Organization has predicted will outpace ischemic heart disease as the number one cause of disease burden by 2030 (Tucci and Moukaddam 2017). Mood disorders, such as MDD and anxiety, are often precipitated by chronic stress. Frequent stress exposure causes adaptation to cortisol release, ultimately altering the HPA axis and stress response (Nemeroff CB 2004; de Kloet et al. 2005; Hammen et al. 2009). Therefore, while complex mood disorders diagnosed with self-report questionnaires (such as MDD) cannot be directly modeled in rodents, chronic stress paradigms can be used to recapitulate some behavioral phenotypes. However, common chronic stress paradigms used in mice, such as corticosterone administration and classic forms of chronic social defeat stress, were developed for males and are not as effective in females. Because of this, most historic pre-clinical chronic stress paradigms exclude females, despite MDD diagnoses occurring twice as frequently in women as in men (Kessler 2003; Kessler et al. 2005; Beery and Zucker 2011). Given the prevalence of MDD in women, there is a need for increased use of female mice in preclinical research. Our lab previously reported a variation of chronic social defeat stress that permits inclusion of female mice (Yohn et al. 2019b).

Chronic social defeat stress (CSDS) is a paradigm in which experimental intruder mice experience repeated stressful bouts of aggression from a resident CD-1 aggressor mouse.

Following these defeat sessions, a social interaction test is used to define susceptible (SUS) mice, which display behavioral and neural changes consistent with increased social avoidance, and resilient (RES) mice, which do not show these changes (Krishnan et al. 2007; Golden et al. 2011). CSDS exploits male rodent territorial aggression. While this behavior is natural in male mice, most strains of female mice are less aggressive, and aggressive male mice tend to mount rather than attack females (Debold and Miczek 1984; Lopez and Bagot 2021). Female resident aggressor mice do not produce the necessary bouts of attack characterizing CSDS unless co-housed with a castrated male (Newman et al. 2019), as female-female aggression in mice is limited to specific territorial species (Trainor et al. 2011; Smith et al. 2013), maternal aggression (Bosch and Neumann 2010; Bourke and Neigh 2012), and following ovariectomy (Debold and Miczek 1984).

There are chronic stress paradigms that work predominately in female mice, such as witness defeat stress and social instability stress; however, these stress paradigms are not as effective in male mice and make direct comparisons between sexes challenging (Haller et al. 1999; Sial et al. 2016; Yohn et al. 2019a; Schuler et al. 2024). Importantly, over the last decade several successful variations of classic social defeat stress were designed to allow inclusion of female mice. These include targeted chemogenetic activation of ventromedial hypothalamus or daily urine applications to increase aggression toward female mice (Takahashi et al. 2017; Harris et al. 2018). In addition, we developed chronic non-discriminatory social defeat stress (CNSDS), which simultaneously exposes a male and female C57BL/6J to an aggressor and is effective in both sexes (Yohn et al. 2019b). CNSDS potentially provides the advantage of males and females having a shared and similar experience during defeat sessions, which could make direct comparisons between sexes more meaningful.

We previously demonstrated that CNSDS yields aggression toward both male and female intruder mice, which resulted in behavioral and neuroendocrine responses consistent with chronic stress exposure. Importantly, we also found that CD-1 aggressor mice attack female intruders far less than male intruders, which has been replicated in many other studies that have utilized or adapted the CNSDS protocol (Yohn et al. 2019b; Zhu et al. 2023; Smith et al. 2023; Bennett et al. 2024). In addition, some labs report that they did not observe increased social avoidance, or susceptibility, in female mice (Parel et al. 2023), which suggests that female mice may display susceptibility and response to chronic stress differently than male mice (Willmore et al. 2022; Balouek et al. 2023; Schuler et al. 2024). Despite these drawbacks, many studies have used CNSDS as an effective paradigm to investigate behavioral and physiological effects of chronic stress in female mice (Yohn et al. 2019b; Dieterich et al. 2021; Zhu et al. 2023; Balouek et al. 2023; Smith et al. 2023; Bennett et al. 2024). In this study, one aim is to update the CNSDS procedure to better facilitate the inclusion of female mice in chronic stress research.

Social interaction test-defined susceptible and resilient phenotypes following CNSDS and other chronic social defeat stress procedures are correlated with changes in sucrose preference and avoidance behaviors (Venzala et al. 2012; Macedo et al. 2018; Yohn et al. 2019b; Guo et al. 2020). However, surprisingly few studies have asked how chronic stress affects more complex reward-related behaviors, especially considering dysfunctional reward processing is a core feature that frequently persists after treatment in mood disorders (Admon and Pizzagalli 2015). A second aim of the study was to determine whether CNSDS-induced susceptible and resilient behavioral phenotypes extend to outcome devaluation, which involves satiety-based devaluation of a food reward during an instrumental operant conditioning task. Importantly, we previously reported that chronic stress exposure abolishes sensitivity to a devalued outcome in females and attenuates sensitivity in male mice (Dieterich et al. 2021).

Here, we introduce an improved screening protocol to identify and rank CD-1 aggressors for CNSDS. We also introduce a new SIT ratio calculation that categorizes susceptibility to CNSDS by directly assessing social avoidance to the CD-1 aggressor rather than general social avoidance, and show that despite receiving fewer attacks, female mice are equally susceptible to CNSDS as male mice. Finally, we investigate the effect of CNSDS on satiety-based outcome devaluation and observe a novel effect of stress susceptibility on sensitivity to a devalued outcome.

## METHODS

### Subjects

Eight-week-old male and female C57BL/6J mice (n=52) were purchased from Jackson Laboratories (Bar Harbor, ME) and retired breeder male CD-1 mice (n=70) were purchased from Charles River Laboratories. Mice were maintained on a 12L:12D schedule with lights turning on at 6 a.m. and turning off at 6 p.m. Throughout the CNSDS procedure, all food and water were provided ad libitum. During the outcome devaluation procedure, mice were food restricted to 85% of their body weight. Behavioral testing took place in the morning, between the hours of 8 a.m. and 1 p.m. All experiments were conducted in compliance with NIH laboratory animal care guidelines and approved by Rutgers University Institutional Animal Care and Use Committee (IACUC).

### CD-1 Aggressor Screening

Retired CD-1 breeder adult males (n=70) were purchased from Charles River Laboratory (Wilmington, MA). To screen for aggressive behavior, screener (non-experimental) female and male C57BL/6J mice aged 8-20 weeks were simultaneously placed in the home cage of a CD-1 mouse (Fig 1A). Latency to attack (in seconds), the number of attacks against male and female intruder mice, the number of bouts of submissive posture to the CD-1 aggressor, and the number of times the CD-1 mounted the intruder mice were recorded. CD-1 mice that attacked the female screener mice within the 5-minute screening sessions were selected for further analysis. An attack was defined as a bite or scratch to the intruder mouse and often included wrestling behavior. CD-1 mice were then ranked by average number of attacks across the three days of screening, and CD-1 mice with the highest average attacks (an average of at least 1 attack per screening session was used as a minimum threshold) were chosen as aggressors.

**Fig 1.**
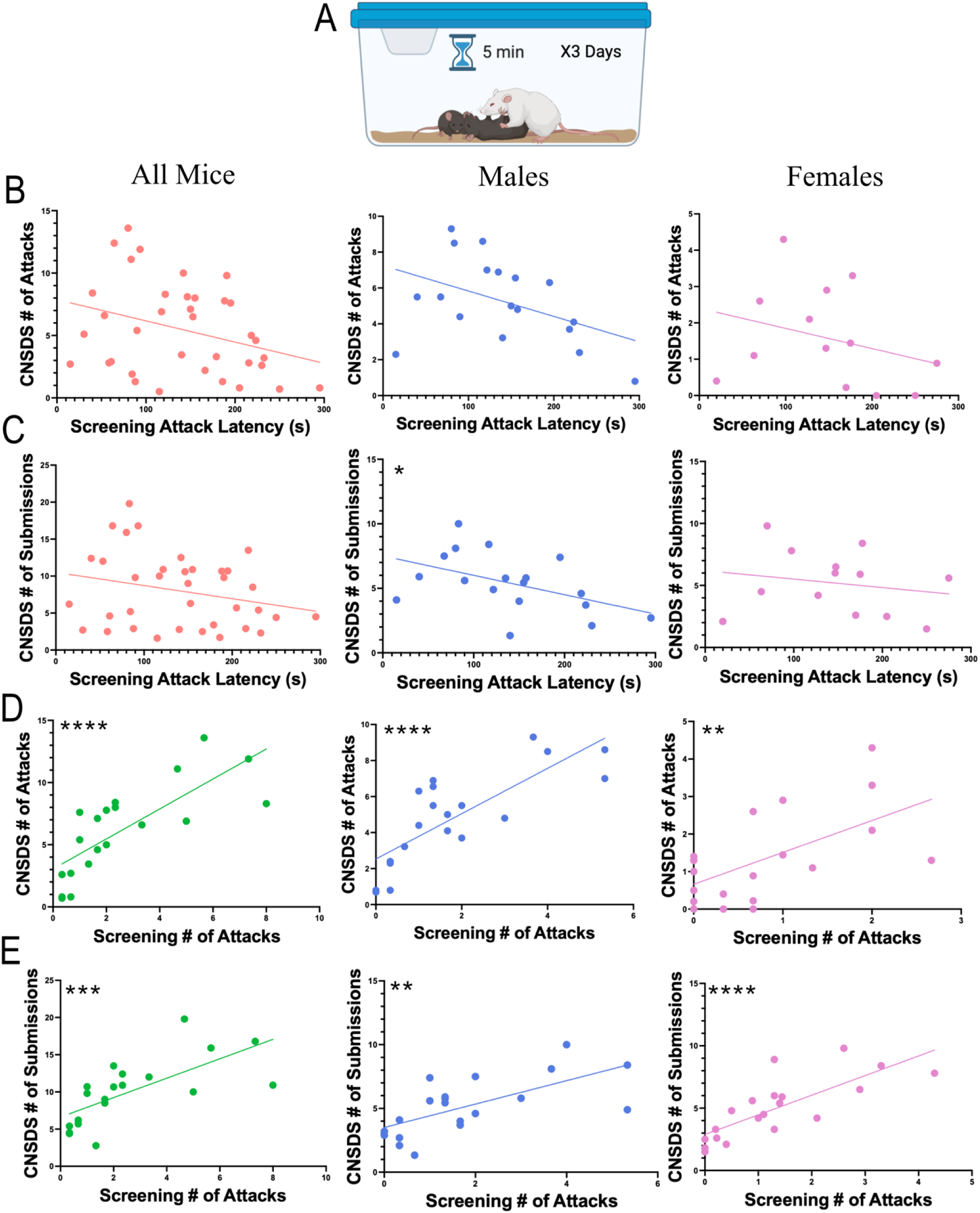
Comparison of CD-1 aggression during screening and CNSDS defeat sessions. A) During screening, CD-1 aggressors are exposed to a male and female C57BL/6J intruder for a 5-minute session for three consecutive days. During this time, number of attacks, latency to first attack, number of submissions, and number of mountings are recorded. B) Linear regressions showing the correlation between the average latency to attack during screening and the average number of attacks per CNSDS session by CD-1 in all mice (R^2^=0.1043, F(1,34)=3.957, p=0.0548), males (R^2^=0.1954, F(1,16)=3.886, p=0.0662), and females (R^2^=0.08856, F(1,11)=1.069, p=0.3234). C) Linear regressions showing the correlation between the average latency to attack during screening and the average number of submissions received per CNSDS session by CD-1 in all mice (R^2^=0.06308, F(1,34)=2.289, p=0.1395), males (R^2^=0.2291, F(1,16)=4.755, p=0.0445), and females (R^2^=0.03748, F(1,11)=0.4284, p=0.5262). D) Linear regressions showing the correlation between the average number of attacks during screening and the average number of attacks per CNSDS session by CD-1 in all mice (R^2^=0.5896, F(1,18)=25.86, p<0.0001), males (R^2^=0.6249, F(1,18)=29.99, p<0.0001), and females (R^2^=0.3496, F(1,18)=9.674, p=0.0060). E) Linear regressions showing the correlation between the average number of attacks during screening and the average number of submissions received per CNSDS session by CD-1 in all mice (R^2^=0.4741, F(1,18)=16.23, p=0.0008), males (R^2^=0.4355, F(1,18)=13.89, p=0.0015), and females (R^2^=0.5971, F(1,18)=26.67, p<0.0001). *p<0.05, **p<0.01, ***p<0.001, ****p<0.0001

### Chronic non-discriminatory social defeat stress

Adult male and female C57BL/6J mice were randomly assigned to either CNSDS or no stress control groups. During the experimental phase, male C57BL/6J mice were matched with the same female C57BL/6J mice throughout the social defeat. The male and female C57BL/6J pairs were placed into the home cage of a novel CD-1 aggressor for 10 consecutive daily 5-min sessions (Fig 2A). C57BL/6J pairs were shifted daily so CD-1 aggressors did not interact with the same intruder pair twice. On alternating days, either the male or female C57BL/6J mouse was co-housed with the CD-1 aggressor with which they had interacted, while the other was co-housed with a novel CD-1. The novel CD-1 was either an aggressor that was not selected for bouts of social defeat or a non-aggressor. Co-housed subjects were separated by clear, perforated plexiglass barriers that permitted sensory, but not physical, interaction. In total, both C57BL/6J male and female mice experienced 5 days of sensory exposure to an aggressor CD-1 they had just interacted with and 5 days of sensory exposure to a different CD-1. During the 5-min sessions, attack latency and frequency, submission behavior, and mounting behavior were measured and recorded for each of the 10 days. The no stress control group of female and male C57BL/6J mice were placed into a cage and allowed to interact for 5 min in the absence of an aggressor CD-1 male over the 10 days. These opposite sex control mice were housed on either side of a perforated, plexiglass divider and placed on a separate rack from chronic stress mice for 24 h until the next interaction session (Fig 2B).

**Fig 2.**
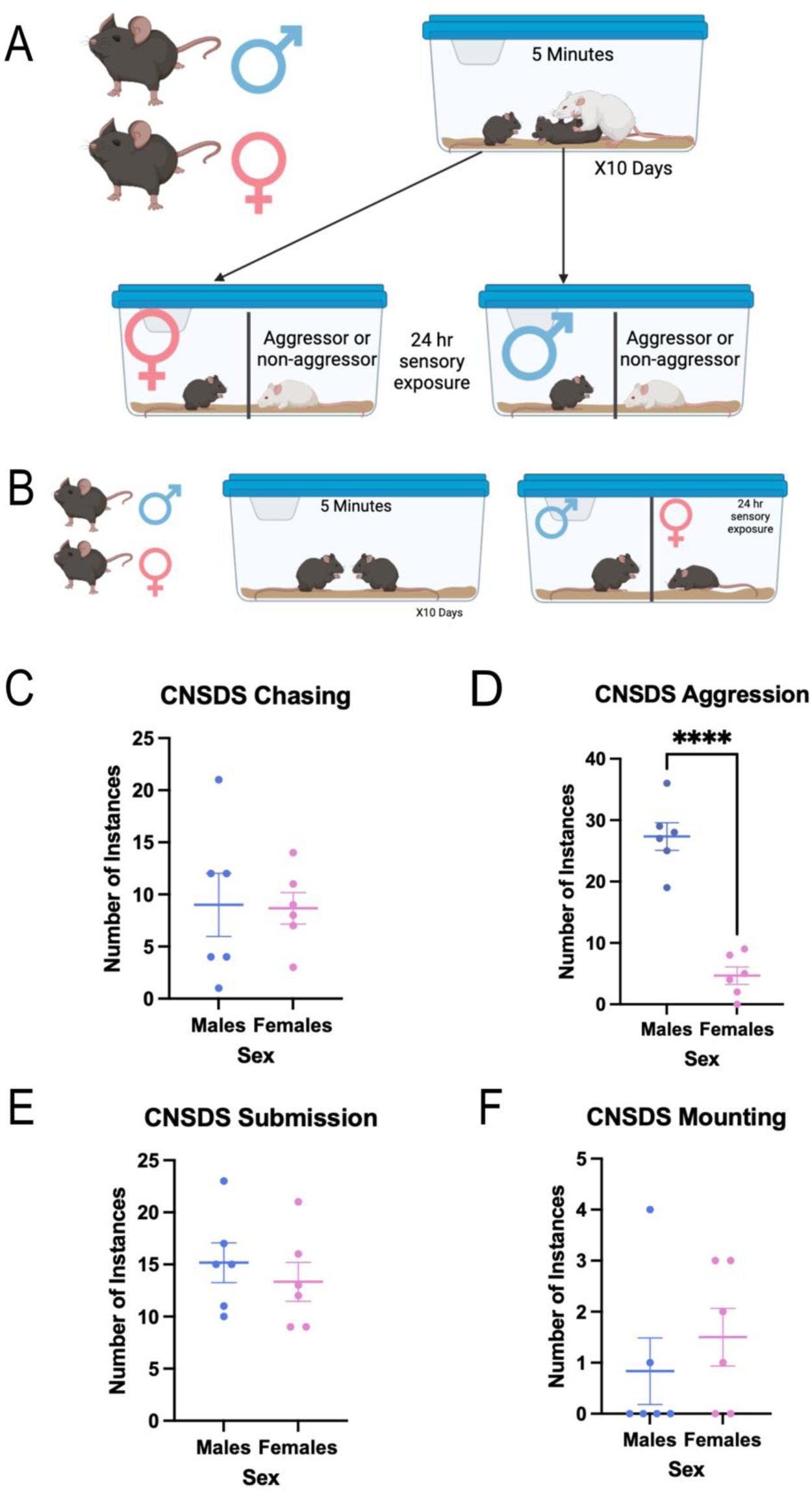
Male intruder mice are attacked significantly more than female mice, but male and female mice show equal instances of submission. A) The male and female C57BL/6J pairs were placed into the home cage of novel CD-1 aggressors for 10 consecutive daily 5-min sessions. B) Control C57BL/6J mice were allowed to interact for 5 min in the absence of an aggressor CD-1 male over the 10 days and were co-housed on either side of a perforated, plexiglass divider. C) There was no significant difference between the number of occurrences of chasing male and female intruders by CD-1 aggressors (t(10)=0.09825, p=0.9237). D) CD-1 aggressor mice attacked male intruders significantly more than female intruders (t(10)=8.513, p<0.0001). E) There was no significant difference between the number of occurrences of male and female submission to CD-1 aggressors (t(10)=0.6862, p=0.5082). F) There was no significant difference between the number of occurrences of mounting male and female intruders by CD-1 aggressors (t(10)=0.7727, p=0.4576). p****<0.0001

### Behavioral Testing

#### Social Interaction Test

To assess susceptibility and resilience status, we ran a social interaction test (SIT) in the presence of a novel CD-1 and a novel C57 mouse. Mice were placed in a plexiglass open field arena (43cm x 43 cm) for three consecutive 2.5-min trials. The first trial had no mouse present in a perforated Plexiglass container within the social interaction zone (14cm x 24 cm). The second trial had a novel CD-1 placed in the perforated container within the social interaction zone (Fig 3A). The third trial had a novel C57BL/6J mouse of the opposite sex within the social interaction zone. Overhead cameras recorded behavior, and Bonsai software (Lopes et al. 2015) measured time spent in the pre-defined interaction zone. Social interaction behavior was measured by time spent in the interaction zone during the second (CD-1) and third (C57) trial, with a social interaction ratio calculated: (interaction time with CD-1 present)/(interaction time with opposite sex C57 present). Mice with an interaction ratio <1 were categorized susceptible (SUS), while mice with an interaction ratio >1 were categorized as resilient (RES) to CNSDS. For comparison, social avoidance behavior was also measured by time spent in the interaction zone during the second (CD-1) and first (no mouse) trials, with a traditional social interaction ratio calculated: (interaction time with CD-1 present)/(interaction time with no mouse present).

**Fig 3.**
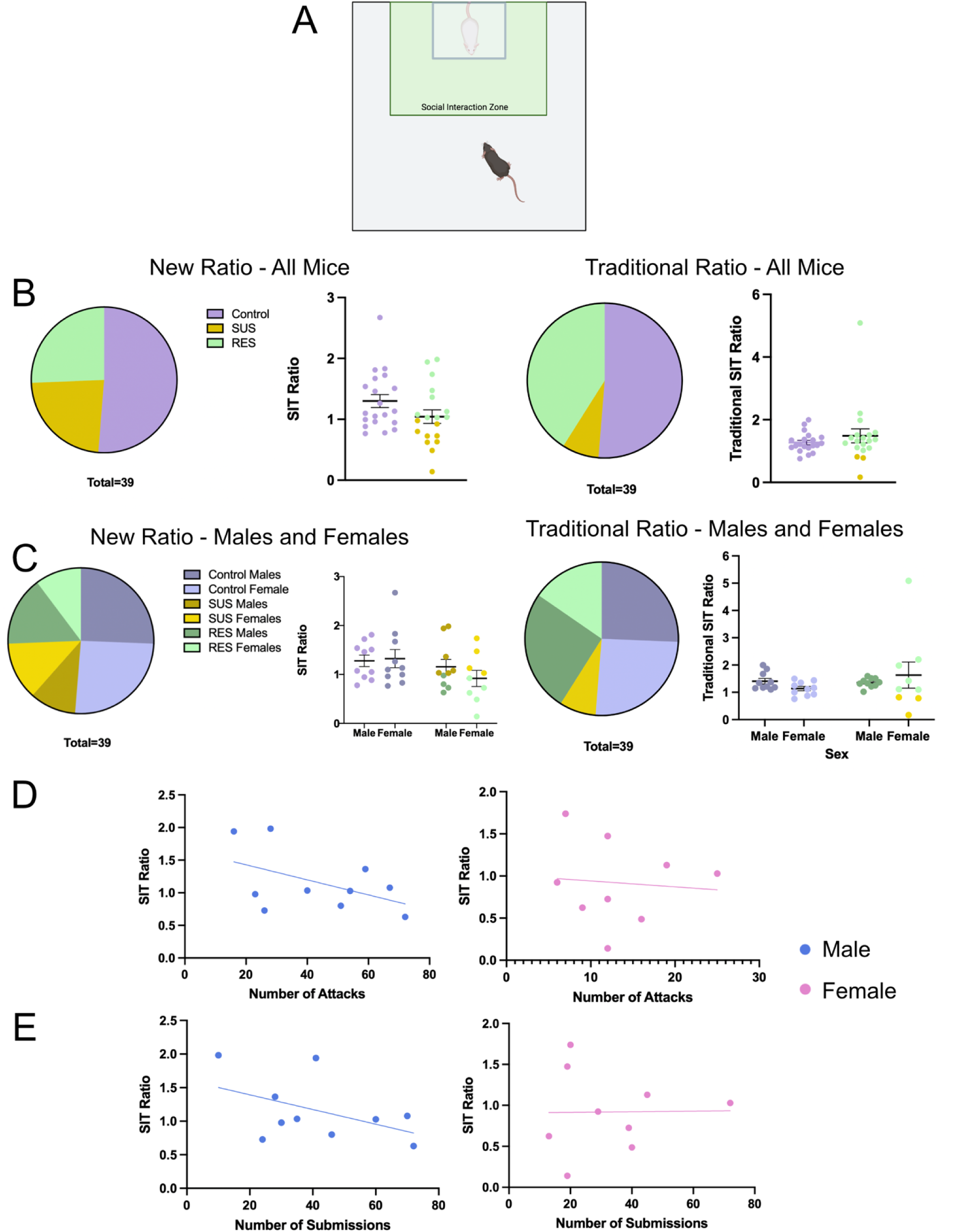
CNSDS results in both susceptible and resilient mice regardless of sex. A) The social interaction test (SIT) was used to analyze social interaction and avoidance behavior in the presence of a novel CD-1 mouse. It consists of 3 2.5-minute trials, the first of which has no mouse present within the social interaction zone, the second has a novel CD-1, and the third has a C57BL/6J of the opposite sex. B) Split of control, SUS, and RES mice after CNSDS using the new SIT ratio (time spent in presence of CD1/time spent in presence of C57) and traditional SIT ratio (time spent in presence of CD1/time spent in absence of all mice). C) Split of control, SUS, and RES mice separated by sex using the new SIT ratio and traditional SIT ratio. D) Linear regressions showing the correlation between the number of attacks received and SIT ratio in males (R^2^=0.2336, F(1,8)=2.438, p=0.1571) and females (R^2^=0.0074, F(1,7)=0.0522, p=0.8258). E) Linear regressions showing the correlation between the number of instances of submissive posture and SIT ratio in males (R^2^=0.2252, F(1,8)=2.326, p=0.1658) and females (R^2^=0.0002, F(1,7)=0.0013, p=0.9718)

#### Outcome devaluation

Male and female C57BL/6J experimental mice were trained to lever press and then tested in standard mouse instrumental conditioning chambers (Med Associates, Fairfax, VT), connected via a power control and interface unit to a dedicated computer with MED-PC IV software (Med Associates, Fairfax, VT) running custom codes. Each instrumental chamber consisted of a single retractable response lever with a reward port delivering 20 mg dustless precision food pellet reinforcers (Bio-Serv, Flemington, NJ), connected by Y-tubing to pellet hoppers. CNSDS-exposed males (n = 10) and females (n = 10) and control males (n=10) and females (n=10) were first exposed to instrumental responding on a Fixed Ratio 1 (FR1) schedule of reinforcement, where each lever press is reinforced by a sucrose pellet. A single lever was ejected into the chamber at the start of each trial, and was retracted following a lever press, coinciding with a single reward pellet being delivered into the reward port for consumption. The lever reappeared after five seconds following a successful trial. Once mice reached the threshold of 20 lever presses per session, they then completed a Variable Ratio 2 (VR2) session, where every 1, 2, or 3 lever presses was reinforced by a sucrose pellet. Once all mice had lever pressed at least 20 times in an FR1 and VR2 session, they progressed to outcome devaluation testing (n=13). Mice that did not reach or surpass the threshold of 20 lever presses were excluded (n=6).

Mice then completed a satiety-based outcome devaluation procedure. Valued and devalued sessions were counterbalanced across two days of testing, separated by a single VR2 session day to re-acquire the response-reward relationship. In the devalued session, mice were pre-fed with reinforcer pellets in the home cage 1 hour before a 5-minute extinction test where the lever was ejected and responses were recorded, but no reinforcers were delivered. In the valued session, mice were pre-fed with standard lab chow in the home cage 1 hour before an identical 5-minute extinction test (Fig 4B). Devalued and valued sessions were counterbalanced between groups, with half of the mice completing the valued session first and the devalued session second, and the other half completing the devalued session first and the valued session second. Response in “devalued’ and “valued” sessions were measured and compared to determine the effect of CNSDS on responding in both sessions.

**Fig 4.**
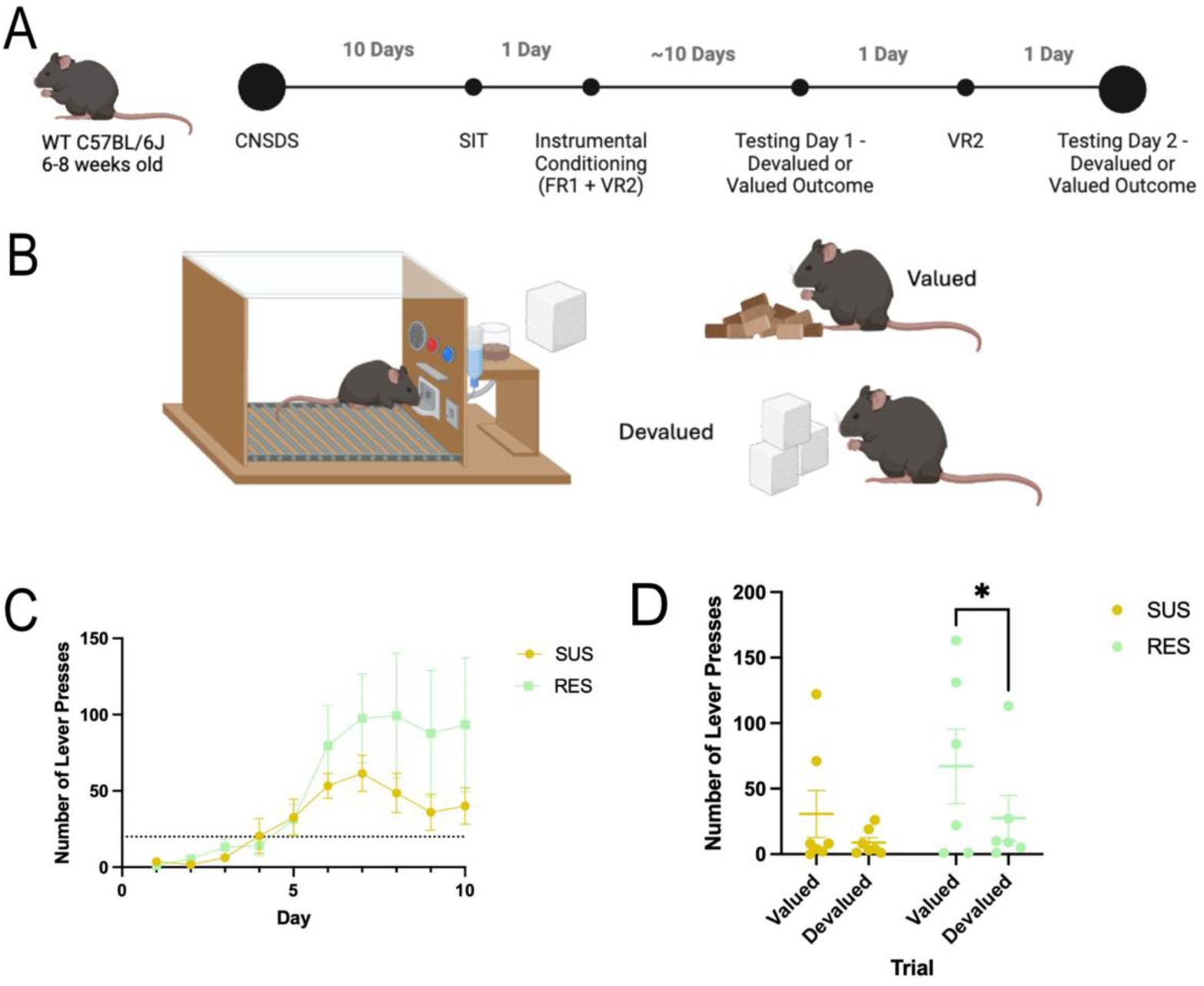
Sensitivity to outcome devaluation is influenced by susceptibility to CNSDS. A) Timeline of CNSDS, operant training, and counterbalanced outcome devaluation testing. B) Mice were pre-fed with either standard lab chow or reinforcer pellets one hour before a 5-minute extinction operant test. C) Training data for the operant task on an FR1 or VR2 schedule. Days 1-6 are FR1, and Days 7-10 are VR2. The dotted line denotes the threshold of 20 lever presses. A 2-way RM ANOVA showed an effect of training day (F(9,108) = 8.633, p=0.0024), but not of susceptibility (F(1,12) = 2.056, p=0.1771) or an interaction between training day and susceptibility (F (9, 108) = 1.320, p=0.2348) on average number of lever presses across the ten days. Planned multiple comparisons for each day show that RES mice do not press significantly more on training days than SUS mice. D) RES mice are more sensitive to satiety-based outcome devaluation than SUS mice. A 2-way RM ANOVA showed an effect of trial (F(1,11) = 7.264, p=0.0208), but not susceptibility F(1,11) = 1.398, P=0.2620) or an interaction between trial and susceptibility (F(1,11) = 0.6006, p=0.4547) on number of lever presses in outcome devaluation. Planned multiple comparisons to assess sensitivity to devaluation in RES and SUS mice show that RES mice press significantly more during the valued trial than the devalued trial (p=0.0375) while SUS mice do not (p=0.1853). *p<0.05

### Statistical Analyses

To investigate the correlation of attack behavior (attack latency and attack frequency) during screening to attack behavior (attack frequency) during CNSDS defeat sessions, a linear regression was used. To assess sex differences in chasing, submission, attack, and mounting frequency, independent samples t-tests were used. To assess the distribution of stress susceptibility compared to the expected outcome, binomial tests were run. To investigate the effect of number of attacks and number of instances of submissive posture on susceptibility to stress, a linear regression was used. To assess the effect of stress susceptibility on outcome devaluation training, a 2 x 2 repeated measures ANOVA with planned Fisher’s LSD comparison test was used. To assess the effect of stress susceptibility on sensitivity to outcome devaluation, a repeated measures 2 x 2 ANOVA with planned Fisher’s LSD comparisons was performed.

GraphPad Prism 10 was used for all analyses.

## RESULTS

### Number of attacks during screening, but not latency to attack, correlates with number of attacks during CNSDS in male and female mice

Published screening parameters for CNSDS and other social defeat protocols state that the aggressor must attack the intruder mice within 60 seconds every day for three days, which is the common threshold in other social defeat stress protocols (Golden et al., 2011; Yohn et al., 2019). An important difference with CNSDS is that aggressors are screened with both male and female intruder mice (Fig. 1A). However, with CNSDS screening many CD-1 mice do not attack within the first 60 seconds, resulting in few mice reaching the threshold to be considered an aggressor despite eventually attacking the intruder mice within the five-minute screening period. It is unclear if latency to attack during CNSDS screening is correlated with increased aggression during CNSDS defeat sessions. Therefore, we performed simple linear regressions to determine if latency to attack and number of attacks during screening significantly correlated with several measures of aggression during CNSDS defeat sessions.

The average CD-1 latency to attack during screening did not significantly correlate to number of attacks per CNSDS defeat session across all mice (R^2^=0.1043, F(1,34)=3.957, p=0.0548).This was consistent for both attacks against male intruders (R^2^=0.1954, F(1,16)=3.886, p=0.0662) and female intruders (R^2^=0.08856, F(1,11)=1.069, p=0.3234) (Fig 1B). Across all mice, the average latency to attack during screening also did not significantly correlate to number of submissions received by CD-1 aggressors per CNSDS defeat session (R^2^=0.06308, F(1,34)=2.289, p=0.1395) (Fig. 1C). While this was consistent for submissions from female intruders (R^2^=0.03748, F(1,11)=0.4284, p=0.5262), the correlation between average latency to attack during screening and the number of submissions received from male intruders was significant (R^2^=0.2291, F(1,16)=4.755, p=0.0445) (Fig 1C).

By contrast, the average number of attacks during screening was significantly correlated to the number of attacks per CNSDS defeat session (R^2^=0.5896, F(1,18)=25.86, p<0.0001), and this was consistent for attacks against male intruders (R^2^=0.6249, F(1,18)=29.99, p<0.0001) and female intruders (R^2^=0.3496, F(1,18)=9.674, p=0.0060) (Fig 1D). The average number of attacks during screening also significantly correlated to the number of submissions received by CD-1 aggressors per CNSDS defeat session (R^2^=0.4741, F(1,18)=16.23, p=0.0008), and this was consistent for submissions from male (R^2^=0.4355, F(1,18)=13.89, p=0.0015) and female (R^2^=0.5971, F(1,18)=26.67, p<0.0001) intruders (Fig 1E). These data suggest that the average number of attacks during screening is a better predictor of CD-1 aggression during CNSDS defeat sessions than latency to attack.

### Male intruder mice are attacked significantly more than female mice, but male and female mice show equal instances of submission

Based on these data, we defined aggressive mice during screening as having attacked the female and averaging at least 1 attack per day. During CNSDS, CD-1 aggressor mice chased male intruders an average of 9 times per defeat session, and chased female mice an average of 8.667 times per session (t(10)=0.09825, p=0.9237) (Fig 2C). CD-1 aggressor mice attacked male intruders an average of 27.33 times per session, while only attacking female intruders an average of 4.667 times per session (t(10)=8.513, p<0.0001) (Fig 2D). Male intruder mice submitted to the CD-1 aggressor an average of 15.17 times per session, while female intruder mice submitted an average of 13.33 times per session (t(10)=0.6862, p=0.5082) (Fig 2E). Male intruder mice were mounted by the CD-1 aggressor an average of 0.8333 times per session, while female intruders were mounted an average of 1.5 times per session (t(10)=0.7727, p=0.4576) (Fig 2F). These data show that male intruder mice are attacked significantly more than female intruder mice during CNSDS defeat sessions, while there are equal instances of submission to, chasing by, and mounting by the CD-1 aggressor in male and female intruder mice.

### CNSDS results in both susceptible and resilient mice regardless of sex

To determine if CNSDS using CD-1 aggressors screened with the new parameters would result in susceptible and resilient mice, we ran the social interaction test (SIT) to analyze social interaction and avoidance behavior in the presence of a CD-1 mouse. The SIT ratio was calculated based on the time spent in the social interaction zone when the CD-1 was present compared to a C57 mouse of the opposite sex being present. Of the 19 mice exposed to stress, 9 were susceptible (SUS) and the other 10 were resilient (RES) (Fig 3B “New Ratio – All Mice”). A binomial test showed that the outcome of SUS vs RES mice was not significantly different from the expected 60/40 split (p=0.3490) (Golden et al. 2011). Of the 10 males exposed to stress, 4 were SUS and 6 were RES. Of the 9 females exposed to stress, 5 were SUS and 4 were RES (Fig 3C “New Ratio – Males and Females”). The traditional SIT ratio was also calculated based on the time spent in the social interaction zone when the CD-1 was present compared to no mouse being present. Of the 19 mice exposed to stress, 3 were SUS and the other 16 were RES (Fig 3B “Traditional Ratio – All Mice”). A binomial test showed that the outcome of SUS vs RES mice was significantly different from the expected 60/40 split (p=0.0002). Of the 10 males exposed to stress, 0 were SUS and 10 were RES. Of the 9 females exposed to stress, 3 were SUS and 6 were RES (Fig 3C “Traditional Ratio – Males and Females”). These data confirm that CNSDS results in susceptible and resilient populations using the novel SIT ratio in both male and female C57BL/6J mice.

### Number of attacks and submissions does not influence susceptibility to stress

To determine if number of attacks received or instances of submissive posture influences susceptibility to stress, we ran linear regressions to investigate the correlation between instances of these behaviors and SIT ratio. Interestingly, the number of attacks received did not correlate to SIT ratio in male (R^2^=0.2336, F(1,8)=2.438, p=0.1571) or female (R^2^=0.0074, F(1,7)=0.0522, p=0.8258) mice (Fig 3D). The number of instances of submissive posture also did not correlate to SIT ratio in male (R^2^=0.2252, F(1,8)=2.326, p=0.1658) or female (R^2^=0.0002, F(1,7)=0.0013, p=0.9718) mice (Fig 3E). These data suggest that the number of attacks received by intruder mice and the number of submissions to the CD-1 aggressor are not relevant measures for determining susceptibility to stress.

### Sensitivity to outcome devaluation is influenced by susceptibility to CNSDS

We next wanted to assess whether CNSDS-induced susceptible and resilient phenotypes correlated with performance in an outcome devaluation task. We trained male and female C57BL/6J mice that had undergone CNSDS to lever press in an operant task on an FR1 and VR2 schedule for a sucrose reward. Once the mice learned the task, they were pre-fed with either lab chow (valued) or sucrose pellets (devalued) in a counterbalanced satiety-based outcome devaluation task (Fig 4A and 4B). Both SUS and RES mice were able to learn the task and passed the threshold of 20 lever presses. A 2-way repeated measures ANOVA showed an effect of training day (F (9, 108) = 8.633, P=0.0024), but not of susceptibility (F (1, 12) = 2.056, P=0.1771), nor an interaction between training day and susceptibility (F (9, 108) = 1.320, P=0.2348) on average number of lever presses across the ten-day period. Planned Fisher’s LSD multiple comparisons within each day show that RES mice do not press significantly more on training days than SUS mice (Fig 4C).

Mice then completed a satiety-based outcome devaluation procedure. Valued and devalued sessions were counterbalanced across two days of testing, separated by a single VR2 session day to re-acquire the response-reward relationship. A 2-way repeated measures ANOVA showed an effect of trial (F(1,11) = 7.264, p=0.0208), but not susceptibility F(1,11) = 1.398, P=0.2620) or an interaction between trial and susceptibility (F(1,11) = 0.6006, p=0.4547) on number of lever presses in outcome devaluation. We next performed planned Fisher’s LSD multiple comparisons to determine sensitivity to outcome devaluation within the SUS and RES groups. Interestingly, RES mice press significantly more during the valued trial than the devalued trial (p=0.0375) while SUS mice do not press significantly more during the valued trial compared to the devalued trial (p=0.1853) (Fig 4D), demonstrating that RES mice were sensitive to satiety-based outcome devaluation while SUS mice were not.

## DISCUSSION

This study aimed to improve the CNSDS protocol previously published by our lab (Yohn et al. 2019b). First, we updated the CD-1 aggressor screening protocol. Traditional CSDS and our published CNSDS parameters state that aggressors must attack the intruders within 1 minute (Golden et al. 2011; Yohn et al. 2019). However, we and others have found that inclusion of female mice may delay initial attacks from the CD-1, resulting in many CD-1 mice not reaching the threshold to be considered an aggressor despite eventually attacking both male and female intruders within the five-minute screening period. We show that latency to attack the male and female intruder does not correlate to aggression displayed during CNSDS, while the number of attacks across the three days of screening is a better predictor of overall aggression. CD-1 mice that attacked an average of 1 or more times during screening attacked at least 3 times during CNSDS (Fig 1D). Therefore, we defined a threshold of 1 attack per screening session on average to predict aggressor behavior. CD-1 mice that do not meet the threshold should be categorized as non-aggressors, while CD-1 mice that meet or surpass the threshold should be categorized as aggressors.

A second aim of this study was to further investigate CNSDS as a valid chronic stress paradigm in both male and female C57BL/6J mice. Our initial report of CNSDS and additional data from other groups using or adapting CNSDS suggest that CD-1 aggressors attack male intruders significantly more than females (Yohn et al, 2019; Zhu et al. 2023; Smith et al. 2023; Bennett et al. 2024). We show that despite receiving fewer attacks (Fig 2D), female mice show similar susceptibility to CNSDS as male mice (Fig 3) when using a new approach to categorization resilience and susceptibility. This new approach compares time spent interacting with a novel CD-1 mouse and time spent interacting with a novel C57 of the opposite-sex, which specifically assesses social avoidance of the aggressor. Decreased social interaction is a common behavioral phenotype observed after social defeat stress (Krishnan et al. 2007); however, some studies report typical social behavior in female mice in response to CNSDS (Parel et al. 2023; Schuler et al. 2024). Other studies have investigated alternative methods of measuring susceptibility to chronic stress in female mice, such as close vigilance (Willmore et al. 2022), change in velocity (Schuler et al. 2024), and novelty-suppressed feeding (Parel et al. 2023).

While highly useful and informative, these measures do not allow for direct comparison between male and female subjects.

Importantly, by defining the social interaction ratio as time spent interacting with a novel CD-1 mouse compared to time spent interacting with a C57 mouse of the opposite sex, we can investigate specific avoidance of the stressor rather than general social avoidance. Traditional chronic social defeat stress paradigms investigate social avoidance by comparing time spent interacting with a novel CD-1 mouse to time spent without a mouse present, typically resulting in a 60/40 split of SUS and RES mice respectively (Golden et al. 2011). When using the traditional SIT ratio after CNSDS, we do not see the expected split of SUS and RES mice (Fig 3). CNSDS utilizes pairs of experimental C57 male and female mice that experience social defeat sessions together over the ten days. This is a different experience for experimental C57 mice than traditional social defeat stress that only uses male mice. Because subjects are exposed to multiple mice during stress, it is important to distinguish between social avoidance of the CD-1 aggressor and avoidance of opposite sex C57 mice. By introducing a novel C57 mouse in the social interaction test, we can more specifically investigate social avoidance of a CD-1 aggressor and utilize this to measure susceptibility to the stress.

Despite receiving fewer attacks than males, female mice show susceptibility to the CNSDS procedure. One potential explanation is that, in addition to direct physical attacks, there may be an additional component of vicarious, or witness, stress experienced by the female intruders. Vicarious social defeat stress is a model in which a mouse witnesses the social defeat of another mouse, without being physically attacked by the aggressor, and can lead to effects such as increased avoidance behaviors, increased corticosterone levels, and decreased sucrose preference (Sial et al. 2016; Iñiguez et al. 2018; Schuler et al. 2024). A recent study by Schuler et al. (2024) investigated both witness social defeat stress and CNSDS and reported that SUS males display decreased SIT ratios while SUS females display reduced changes in velocity. These similarities observed by Schuler et al in females across CNSDS and witness defeat stress could suggest that there are similar stress components or experiences across both procedures.

A third aim of this study was to investigate whether CNSDS-induced susceptible and resilient phenotypes correlated with reward valuation behavior. Importantly, reward valuation behavior is highly conserved across species (Der-Avakian et al. 2016), and abnormal reward valuation is observed in major depressive disorder (Rupprechter et al. 2021). Therefore, assessing chronic stress effects on reward valuation has a high translational value. Chronic stress disrupts brain systems involved in reward processing, leading to altered decision making and diminished sensitivity to rewards (Ironside et al. 2018). We previously reported that chronic stress results in the abolishment of sensitivity to satiety-based outcome devaluation, leading to similar responding in the valued and devalued trials (Dieterich et al. 2021). Importantly, this is the first study to investigate whether susceptible and resilient phenotypes to chronic stress correlate with sensitivity to outcome devaluation. Resilient mice pressed significantly more in the valued task compared to the devalued task, while susceptible mice pressed less in both valued and devalued trials (Fig 4). This suggests that while susceptible mice have an abolished or decreased sensitivity to satiety-based outcome devaluation (as expected with chronic stress), resilient mice are more sensitive. One potential explanation is an overall decrease of responding specifically in the susceptible group. Therefore, it is possible that susceptible mice are less likely to expend effort in an operant task (Dieterich et al. 2021). A potential future experiment is to perform a progressive ratio or concurrent choice operant task after mice are exposed to CNSDS and SIT to determine if susceptible mice show a decreased breakpoint and less lever pressing than resilient mice. Another future experiment could be to determine if stress susceptibility impacts motivation behaviors that are affected by CNSDS, such as effort-related choice (Dieterich et al. 2021). Mice categorized into different stress susceptibility groups may show changes in willingness to exert effort, which could support our findings from the outcome devaluation task.

There is a growing need for preclinical research using female mice to study translationally relevant behaviors for mood disorders. Our updated CNSDS paradigm will facilitate the inclusion of female mice in preclinical chronic stress research and better permit direct comparisons between males and females. CNSDS is effective in both sexes and allows increased inclusion of females in the preclinical study of neurobiological circuits involved in mood disorders such as MDD. In addition, CNSDS allows for investigation of the mechanisms underlying stress susceptibility and resilience in both sexes. Ultimately, CNSDS should allow for the study of whether new therapeutic interventions for mood disorders may be effective in both men and women.

## Acknowledgements

The authors declare no conflict of interest. This research was funded by National Institute of Mental Health Grant R01MH123544 (BAS) and the IRACDA at Rutgers: INSPIRE Postdoctoral Training Program K12GM093854 (KD).

## References

1. Admon R, Pizzagalli DA (2015) Dysfunctional reward processing in depression. Curr Opin Psychol 4:114–118. 10.1016/j.copsyc.2014.12.011

2. Balouek J-A, Mclain CA, Minerva AR, et al (2023) Reactivation of Early-life stress-sensitive neuronal ensembles contributes to lifelong stress hypersensitivity. J Neurosci 43:5996– 6009. 10.1523/JNEUROSCI.0016-23.2023

3. Beery AK, Zucker I (2011) Sex bias in neuroscience and biomedical research. Neurosci Biobehav Rev 35:565–572. 10.1016/j.neubiorev.2010.07.002

4. Bennett SN, Chang AB, Rogers FD, et al (2024) Thyroid hormones mediate the impact of early-life stress on ventral tegmental area gene expression and behavior. Horm Behav 159:105472. 10.1016/j.yhbeh.2023.105472

5. Bosch OJ, Neumann ID (2010) Vasopressin released within the central amygdala promotes maternal aggression. Eur J Neurosci 31:883–891. 10.1111/j.1460-9568.2010.07115.x

6. Bourke CH, Neigh GN (2012) Exposure to repeated maternal aggression induces depressive-like behavior and increases startle in adult female rats. Behav Brain Res 227:270–275. 10.1016/j.bbr.2011.11.001

7. de Kloet ER, Joëls M, Holsboer F (2005) Stress and the brain: from adaptation to disease. Nat Rev Neurosci 6:463–475. 10.1038/nrn1683

8. Debold J, Miczek K (1984) Aggression persists after ovariectomy in female rats. Horm Behav 18:177–190. 10.1016/0018-506X(84)90041-2

9. Der-Avakian A, Barnes SA, Markou A, Pizzagalli DA (2016) Translational Assessment of Reward and Motivational Deficits in Psychiatric Disorders. In: Robbins TW, Sahakian BJ (eds) Translational Neuropsychopharmacology. Springer International Publishing, Cham, pp 231–262

10. Dieterich A, Liu T, Samuels BA (2021) Chronic non-discriminatory social defeat stress reduces effort-related motivated behaviors in male and female mice. Transl Psychiatry 11:125. 10.1038/s41398-021-01250-9

11. Golden SA, Covington HE 3rd, Berton O, Russo SJ (2011) A standardized protocol for repeated social defeat stress in mice. Nat Protoc 6:1183–1191. 10.1038/nprot.2011.361

12. Guo Q, Wang L, Yuan W, et al (2020) Different effects of chronic social defeat on social behavior and the brain CRF system in adult male C57 mice with different susceptibilities. Behav Brain Res 384:112553. 10.1016/j.bbr.2020.112553

13. Haller J, Fuchs E, Halász J, Makara GB (1999) Defeat is a major stressor in males while social instability is stressful mainly in females: towards the development of a social stress model in female rats. Brain Res Bull 50:33–39. 10.1016/s0361-9230(99)00087-8

14. Hammen C, Kim EY, Eberhart NK, Brennan PA (2009) Chronic and acute stress and the prediction of major depression in women. Depress Anxiety 26:718–723. 10.1002/da.20571

15. Harris AZ, Atsak P, Bretton ZH, et al (2018) A Novel Method for Chronic Social Defeat Stress in Female Mice. Neuropsychopharmacology 43:1276–1283. 10.1038/npp.2017.259

16. Iñiguez SD, Flores-Ramirez FJ, Riggs LM, et al (2018) Vicarious social defeat stress induces depression-related outcomes in female mice. Biol Psychiatry 83:9–17. 10.1016/j.biopsych.2017.07.014

17. Ironside M, Kumar P, Kang M-S, Pizzagalli DA (2018) Brain mechanisms mediating effects of stress on reward sensitivity. Curr Opin Behav Sci 22:106–113. 10.1016/j.cobeha.2018.01.016

18. Kessler RC (2003) Epidemiology of women and depression. J Affect Disord 74:5–13. 10.1016/s0165-0327(02)00426-3

19. Kessler RC, Berglund P, Demler O, et al (2005) Lifetime prevalence and age-of-onset distributions of DSM-IV disorders in the National Comorbidity Survey Replication. Arch Gen Psychiatry 62:593–602. 10.1001/archpsyc.62.6.593

20. Krishnan V, Han M-H, Graham DL, et al (2007) Molecular Adaptations Underlying Susceptibility and Resistance to Social Defeat in Brain Reward Regions. Cell 131:391–404. 10.1016/j.cell.2007.09.018

21. Lopes G, Bonacchi N, Frazão J, et al (2015) Bonsai: an event-based framework for processing and controlling data streams. Front Neuroinform 9:7. 10.3389/fninf.2015.00007

22. Lopez J, Bagot RC (2021) Defining valid chronic stress models for depression with female rodents. Biol Psychiatry 90:226–235. 10.1016/j.biopsych.2021.03.010

23. Macedo GC, Morita GM, Domingues LP, et al (2018) Consequences of continuous social defeat stress on anxiety- and depressive-like behaviors and ethanol reward in mice. Horm Behav 97:154–161. 10.1016/j.yhbeh.2017.10.007

24. Nemeroff CB CB (2004) Early-life adversity, CRF dysregulation, and vulnerability to mood and anxiety disorders. Psychopharmacol Bull 38:14–20

25. Newman EL, Covington HE 3rd, Suh J, et al (2019) Fighting females: Neural and behavioral consequences of social defeat stress in female mice. Biol Psychiatry 86:657–668. 10.1016/j.biopsych.2019.05.005

26. Parel ST, Bennett SN, Cheng CJ, et al (2023) Transcriptional signatures of early-life stress and antidepressant treatment efficacy. Proc Natl Acad Sci U S A 120:e2305776120. 10.1073/pnas.2305776120

27. Rupprechter S, Stankevicius A, Huys QJM, et al (2021) Abnormal reward valuation and event-related connectivity in unmedicated major depressive disorder. Psychol Med 51:795–803. 10.1017/S0033291719003799

28. Schuler H, Eid RS, Wu S, et al (2024) Data-driven analysis identifies novel modulation of social behavior in female mice witnessing chronic social defeat stress. Biol Psychiatry. 10.1016/j.biopsych.2024.11.017

29. Sial OK, Warren BL, Alcantara LF, et al (2016) Vicarious social defeat stress: Bridging the gap between physical and emotional stress. J Neurosci Methods 258:94–103. 10.1016/j.jneumeth.2015.10.012

30. Smith A, Hyland L, Al-Ansari H, et al (2023) Metabolic, neuroendocrine and behavioral effects of social defeat in male and female mice using the chronic non-discriminatory social defeat stress model. Horm Behav 155:105412. 10.1016/j.yhbeh.2023.105412

31. Smith AS, Lieberwirth C, Wang Z (2013) Behavioral and physiological responses of female prairie voles (Microtus ochrogaster) to various stressful conditions. Stress 16:531–539. 10.3109/10253890.2013.794449

32. Takahashi A, Chung J-R, Zhang S, et al (2017) Establishment of a repeated social defeat stress model in female mice. Sci Rep 7:12838. 10.1038/s41598-017-12811-8

33. Trainor BC, Pride MC, Villalon Landeros R, et al (2011) Sex differences in social interaction behavior following social defeat stress in the monogamous California mouse (Peromyscus californicus). PLoS One 6:e17405. 10.1371/journal.pone.0017405

34. Tucci V, Moukaddam N (2017) We are the hollow men: The worldwide epidemic of mental illness, psychiatric and behavioral emergencies, and its impact on patients and providers. J Emerg Trauma Shock 10:4–6. 10.4103/0974-2700.199517

35. Venzala E, García-García AL, Elizalde N, et al (2012) Chronic social defeat stress model: behavioral features, antidepressant action, and interaction with biological risk factors. Psychopharmacology (Berl) 224:313–325. 10.1007/s00213-012-2754-5

36. Willmore L, Cameron CM, Yang J, et al (2022) Behavioural and dopaminergic signatures of resilience. Nature 611:124–132. 10.1038/s41586-022-05328-2

37. Yohn CN, Ashamalla SA, Bokka L, et al (2019a) Social instability is an effective chronic stress paradigm for both male and female mice. Neuropharmacology 160:107780. 10.1016/j.neuropharm.2019.107780

38. Yohn CN, Dieterich A, Bazer AS, et al (2019b) Chronic non-discriminatory social defeat is an effective chronic stress paradigm for both male and female mice. Neuropsychopharmacology 44:2220–2229. 10.1038/s41386-019-0520-7

39. Zhu X, Sakamoto S, Ishii C, et al (2023) Dectin-1 signaling on colonic γδ T cells promotes psychosocial stress responses. Nat Immunol 24:625–636. 10.1038/s41590-023-01447-8

